# M&Ms: A software for building realistic Microbial Mock communities

**DOI:** 10.1101/2021.04.21.440404

**Authors:** Natalia García-García, Javier Tamames, Fernando Puente-Sánchez

## Abstract

**Motivation:** Advances in sequencing technologies have triggered the development of many bioinformatic tools aimed to analyze these data. As these tools need to be tested, it is important to simulate datasets that resemble realistic conditions. Although there is a large amount of software dedicated to produce reads from ‘in silico’ microbial communities, often the simulated data diverge widely from real situations.

**Results:** Here, we introduce M&Ms, a user-friendly open-source bioinformatic tool to produce realistic amplicon datasets from reference sequences, based on pragmatic ecological parameters. This tool creates sequence libraries for ‘in silico’ microbial communities with user-controlled richness, evenness, microdiversity, and source environment. M&Ms allows the user to generate simple to complex read datasets based on real parameters that can be used in developing bioinformatic software or in benchmarking current tools. M&Ms also provides additional figures and files with extensive details on how each synthetic community is composed, so that users can make informed choices when designing their benchmarking pipelines.

**Availability:** The source code of M&Ms is freely available from https://github.com/ggnatalia/MMs

## 1 Introduction

Microorganisms dwell every habitable environment in the planet’s biosphere where they play essential roles in sustaining life. Microbiome studies have sharply increased in the last decades, due in part to a parallel development of sequencing technologies. First studies were performed in extreme environments such as acid mine drainages, where the microbial diversity was low; since then, more complex environments such as gut, marine or soils have been studied (White RA *et al.*, 2016). These microbial communities have different levels of complexity, containing from hundreds to thousands of distinct taxa. Species have been traditionally considered the most significant units in microbial ecology (Shappiro & Polz, 2014) but, lately, individuals classified within the same species have been involved in different ecological tradeoffs or in different niches (Rasko *et al.*, 2008, Eren *et al.*, 2013, Kashtan *et al.*, 2014, García-García *et al.*, 2019). Thus, in the last decade, the way of study microbial communities has evolved and factors such as microdiversity, referred to the structural and functional diversity below the species level (Schloter *et al*., 2000), are increasingly relevant.

The most significant technologies for inspecting the profile and function of microbial communities are based on DNA sequencing, like metagenomics and metatranscriptomics. To analyze such amount of data, sophisticated bioinformatic tools have emerged. These new computational methods must be evaluated to ensure a proper functioning. This allows developers to benchmark and compare their software (Baxter *et al.*, 2006, Mangul *et al.*, 2019, Weber *et al.*, 2019). One way to assess the performance of the different algorithms is using simulated data generated on the computer (Engle *et al.*, 1993, Alosaimi *et al.*, 2020), because often appropriate, real data, are not available. However, choosing an artificial dataset that guarantees an optimal performance is not trivial: the simulated data can be biased by the algorithms, cannot capture true experimental variability or can be less complex than real data (Mangul *et al.*, 2019).

There is a large amount of sequencing simulators such as MetaSIM (Richter *et al.*, 2008), Grinder (Angly *et al.*, 2012), insilicoSeqs (Gourlé *et al.*, 2019), PBSIM2 (Ono *et al.*, 2020), SimuSCoP (Yu *et al.*, 2020) (for a thorough review see ref. Alosaimi *et al.*, 2020). These tools are specially designed for simulating read-level artifacts related with sequencing runs or read quality. Their aim is to mimic DNA sequences according to the different sequencing platforms, but not all of them are prepared to simulate realistic microbial communities, or to emulate microbial population heterogeneity.

Other tools are prepared to simulate the dynamics of realistic microbial communities, but they are focused on providing a synthetic 16S rDNA-seq count table that resembles the structure of a microbial community under fluctuating environmental conditions. These tools do not produce simulated sequencing data. Some examples are: the Community Simulator (Marsland *et al.*, 2020) whose aim is simulating microbial population dynamics including environmental conditions throughout time; metaSPARSIM (Patuzzi *et al.*, 2019), a software to generate count matrices resembling real 16S rDNA-seq data; or SPIEC-EASI (Kurz *et al.*, 2015) a sophisticated synthetic microbiome data generator with controllable underlying species interaction topology.

To our knowledge, the first tool designed to produce simulated data from realistic microbial communities is CAMISIM (Fritz *et al.* 2019). CAMISIM produces automatically metagenomic samples emulating different microbial abundance profiles, multi-sample time series, and differential abundance studies (Fritz *et al.* 2019). It also includes real and simulated strain-level diversity, and generates sequencing data using taxonomic profiles, or completeley de novo (Fritz *et al.* 2019). However, CAMISIM does not allow to control community features such as diversity or correlations between taxa.

Selecting the most appropriate bioinformatic tool can be challenging (Mangul *et al.*, 2019) and depends to a large extent on the objectives. The performance tests can contribute the most to the development process and eventually consolidate the tool to handle pragmatic situations (Baxter *et al.*, 2006, Weber *et al.*, 2019). One alternative is the use of the gold standard datasets, usually composed of most known taxa in well characterized environments, but these can hinder the genuine performance of the different tools and lead to overfitting (Mangul *et al.*, 2019).

Here, we describe M&Ms, a user-friendly tool originally written to generate from simple to complex simulated 16S rDNA reads from metagenomic datasets based on desired microbial community’s characteristics such as the microdiversity.

## 2 Methods

M&Ms models a multi-sample microbial abundance profile, includes simulated microdiversity, and generates sequencing data from taxonomic profiles or without them using InSilicoSeqs (Gourlé *et al.*, 2019). M&Ms aims to produce ecologically meaningful artificial microbial communities by means of an abundance profile based on the evenness and richness of real microbial communities, as well as its microdiversity. It also allows for most frequent taxa per environment thanks to a previous study (Tamames *et al.*, 2016). The software produces a FASTQ file and an abundance profile per sample. Also, it produces a FASTQ file that collects the reads of all the samples and a a mothur-formatted groups file with the name of the reads and the sample they belong to.

Simulation with M&Ms has four stages (**Figure 1**):

1. Selection of the community members
2. Microdiversity simulation
3. Microbial abundance distribution assignment
4. Sequencing data simulation using InSilicoSeqs (Gourlé *et al.*, 2019) to produce realistic Illumina reads.

**Figure 1:**
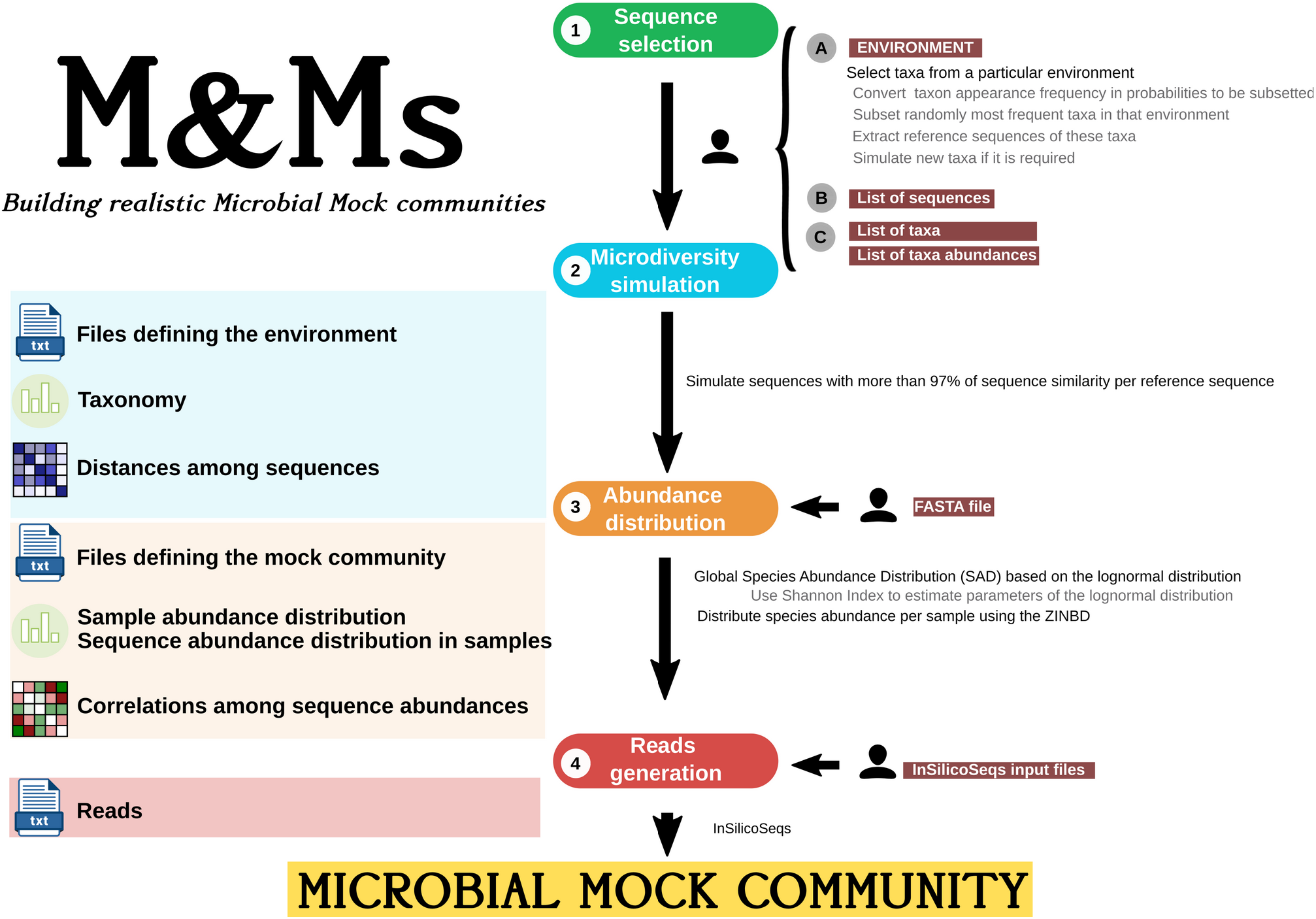
M&Ms pipeline

The software accepts one of the following five independent types of inputs:

- A list of sequences from a particular DB e.g. SILVA database (Yilmaz *et al.*, 2014) from which M&Ms generates a FASTA file, an abundance profile and the simulated sequencing data.
- A list of taxa from the reference database (and optionally their abundances).
- A pre-existing FASTA file with the desired sequences, on the basis of which M&Ms produces the simulated abundance profile and the data.
- A list of 16S rRNA gene sequences and their abundances, so that only simulated sequencing data will be generated.
- A particular environment, so most of the taxa will be selected from that environment, according to a reference table with most frequent taxa (at the genus level) per environment. M&Ms will produce a FASTA file with sequences from species from that environment, an abundance profile and the simulated sequencing data.

To select taxa from a particular environment, M&Ms uses a genus per environment matrix derived in a previous work (Tamames *et al.*, 2016)(see **Supplementary Note 1** for details). For a given pool of possible reference sequences, M&Ms simulates its corresponding microdiversity introducing point mutations.

We use a log-normal distribution to predict the global species abundance distribution (SAD) of the artificial community. Although, since the 1930s, ecologists have developed different models to predict the SAD (McGuill, 2010), those based on the log-normal are considered to be standards to test new models (Shoemaker *et al,* 2017). The lognormal is characterized by a right-skewed frequency distribution that becomes approximately normal under log-transformation. This model reflects the ‘rare biosphere’ (i.e. the fact that most species in a given sample will be present in very low abundances), which is one of the most intensively studied patterns of microbial diversity (Pedrós Alió, 2012; Shoemaker *et al.*, 2017).

The probability density function of the log normal distribution is defined by the mean μ and standard deviation σ, which affects the shape of the distribution. Longuet-Higgins and also Edden (Longuet-Higgins, 1971; Edden, 1971) studied the relation between species diversity and the log-normal distribution of individuals among species. Both proposed the use of the logarithmic standard deviation, σ, as an indicator of the unevenness of species distribution. In particular, σ and the number of species correlates well with the values given by the Shannon and Weaver diversity index (Longuet-Higgins 1971; Edden 1971). The Shannon index (H) is an index commonly used to characterize species diversity in a community that accounts for both abundance and evenness of the species present (Shannon 1948, Hill *et al.*, 2002). We calculate the standard deviation of the log-normal distribution from a specific Shannon Index to approximate the SAD of the mock community, thus the species abundance distribution is more realistic and allow M&Ms to design ecologically meaningful mock communities unlike other equivalent tools. A more detailed explanation of this approximation between Shannon diversity index and the standard deviation of a log-normal distribution can be found in **Supplementary Note 2**.

We then determine the abundance distribution of the different simulated individuals using the Zero-Inflated Negative Binomial Distribution (ZINBD). We use the NORmal-To-Anything (NORTA) approach, which is an approximate technique to generate arbitrary continuous and discrete multivariate distributions from a target correlation matrix, using a multivariate normal (Yahav & Shmuelli, 2011; Kurz *et al.*, 2015). First, we sample from a multivariate normal distribution with zero mean and standard deviation one using a randomly created correlation matrix. For each probability, the Normal cumulative distribution function (CDF) is transformed to the target distribution, in this case, the ZINBD via its inverse CDF (Yahav & Shmuelli, 2011; Kurz *et al.*, 2015). The target distribution is the ZINBD which has been observed to fit properly microbiome data, characterized by having an increasing number of zeros at lower taxonomic levels (Xu *et al.*, 2015; Xia & Sun, 2017).

Finally, M&Ms runs InSilicoSeqs (Gourlé *et al.*, 2019) to simulate metagenomic Illumina sequencing samples with the designed abundance profiles.

## 3 Results

M&Ms has been designed to produce artificial communities, which can be used to test new software in different stages of development and benchmarking. We envisage three different situations: simple mock communities with few sequences (useful in initial stages of software development), complex artificial communities but just one or more samples, convenient in intermediate steps of software development. Additionally, it also facilitates the option of working with the same mock community but with different simulated sequencing parameters in a straightforward way.

M&Ms provides also plots and extra files to effortlessly visualize the main characteristics of the mock community. With all this information, the user can handily establish comparisons among the initial composition of the community and the results of applying any tool. Also, M&Ms have been develop to be flexible, so adding extra improvements such as a better algorithm to simulate microdiversity can be effortlessly implemented.

Therefore, the main advantage of M&Ms consists of automatically obtaining realistic microbial mocks with ease. A comparison with similar tools that apart from generating DNA data, are able to simulate microorganisms distribution is displayed in **Supplementary Note 3**.

