## Supplementary for "M&Ms: A software for building realistic Microbial Mock communities"

### Note 1: Selecting taxa from a specific environment

As it has been mentioned in the main text, we use a genera per environment matrix (Tamames *et al.,* 2016) to select those genera which are more likely to appear in a particular environment (**Table S1**). We assume that genera which are present in more samples from an environment are more relatively abundant in it. According to that, for each environment, we convert these frequencies of occurrence into probabilities of those taxa to be present in that environment normalizing those values by the total counts. For those genera which have not been reported in an environment (they have been observed 0 times in it), we ascribe them a minimum frequency of occurrence (0.005), to account for the possibility that they are rare taxa truly present in the environment, but not detected in the study of (Tamames *et al.,* 2016). For the pool of possible reference sequences in the SILVA database (Yilmaz *et al*., 2014) (classified taxonomically at the genus level), M&Ms carried out a weighted random sampling of 10,000 sequences. In the case that more sequences from a specific taxon are needed, simulated sequences will be created at the species level.

| Environment | | Samples |
|---|---|---|
| Aquatic | Freshwater sediment | 299 |
| | Freshwater saline waters interfase | 88 |
| | Freshwaters | 1999 |
| | Saline waters | 2783 |
| | Soil Freshwaters interfase | 143 |
| | Soil Saline waters interfase | 78 |
| Host associated & Organic | Animal host | 89 |
| | Gut | 1618 |
| | Oral | 90 |
| | Organic | 153 |
| | Other tissue | 248 |
| | Vagina | 22 |
| Terrestrial | Plants | 167 |
| | Saline soil | 67 |
| | Soil | 2549 |
| Thermal | Geothermal | 268 |
| | Hydrothermal | 191 |
| Other | Aerial | 87 |
| | Artificial | 459 |
| | Oil | 202 |

**Table S1. Available environments.** The genera per environment matrix includes 20 different environments and 636 different taxa classified at the genus level.

We use the sequence similarity in the 16S rRNA sequence as the criteria to define species: two sequences with more than a 97% of sequence similarity are considered to belong to the same species (Yarza *et al*., 2014, Roselló-Mora & Amann, 2015). From that pool of 10,000 sequences (representing 10,000 species), M&Ms consecutively subsets one sequence (species) and simulates its corresponding microdiversity until reaching the number of unique sequences required by the user including species and their microdiversity.

This process results in a synthetic community that closely matches the taxon frequencies of the selected environment (**Fig. S1**) and the microdiversity ratio per species required by the user (**Fig. S2**).

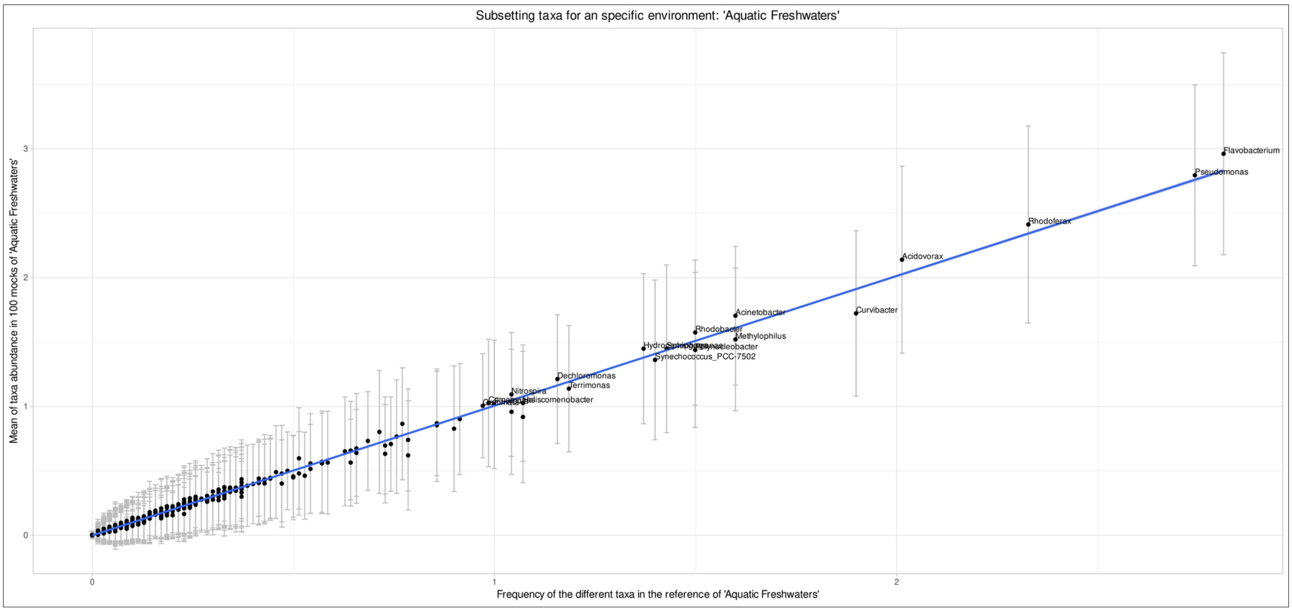


**Fig. S1. M&Ms selects taxa from a specific environment.** Average abundance of selected taxa in 100 mock communities from the 'Aquatic Freshwaters' environment. Most repeated taxa in these mocks are most frequent taxa in that environment according to (Tamames *et al*., 2016). Significant linear regression (pval < 0.001) is shown with a blue line.

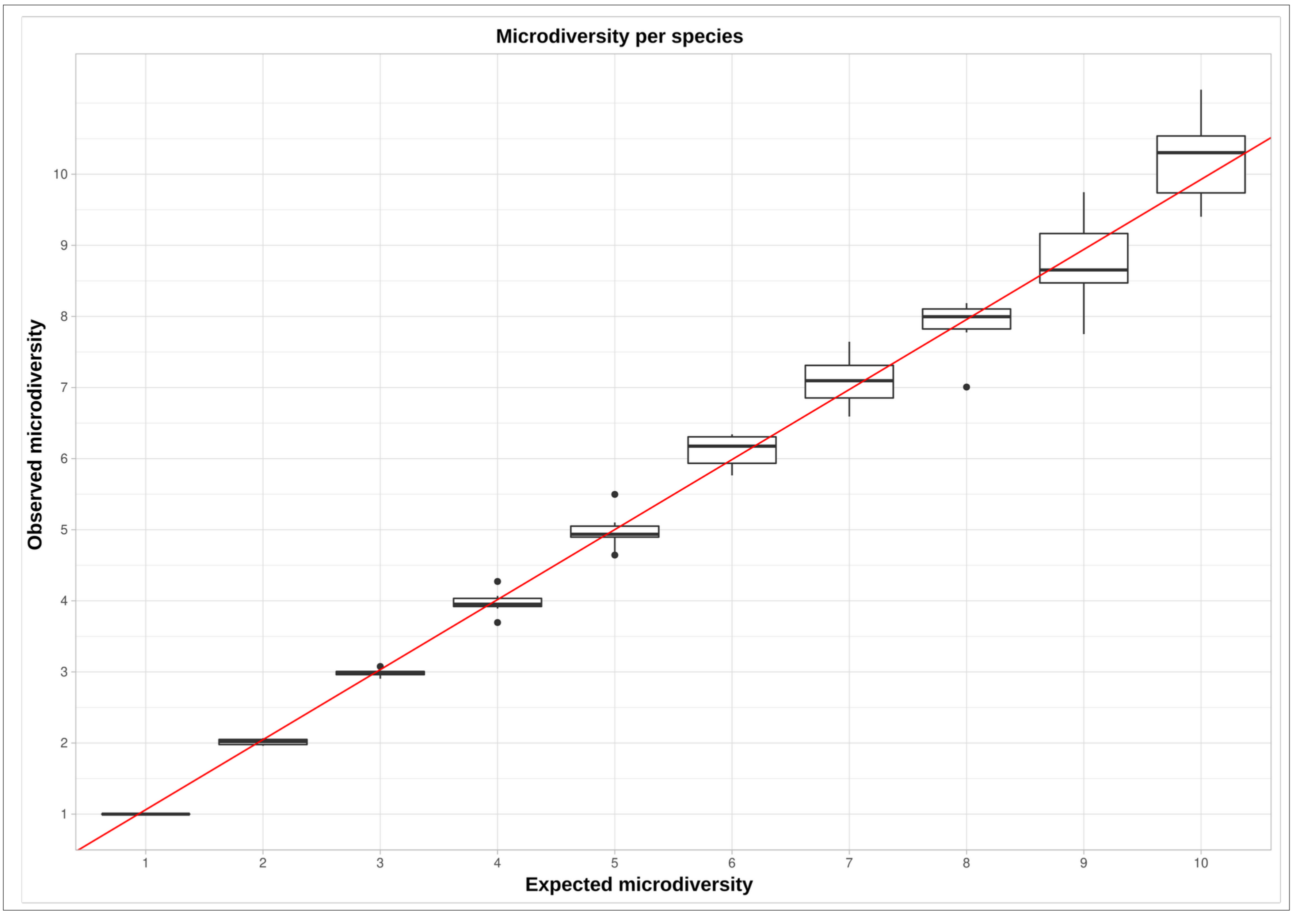


**Fig. S2. M&Ms simulates microdiversity.** Boxplots of the average microdiversity per species in 10 mock communities for different requested values of microdiversity per species. Significant linear regression (pval < 0.001) is shown with a red line.

### Note 2: Shannon Index and the log-normal distribution

One of the main key strengths of M&Ms is the ability of generating artificial microbial communities with a given Shannon diversity index. The Shannon index (H) is an index commonly used to characterize species diversity in a community that accounts for both abundance and evenness of the species present (Shannon 1948, Hill *et al*., 2003):

$$H = -\sum^{S} p_i \cdot \ln(p_i)$$

where:

- H is the Shannon's diversity index
- S is the total number of species in the community (richness)
- $p_i$ is the proportion of S made up of the *-ith* species

In 1971, Longuet and Higgins studied the connection of the log-normal distribution, defined by the mean μ and standard deviation σ, and the evenness of species distribution. They noticed that σ and the number of species correlate well with the values given by the Shannon and Weaver diversity index through the formula:

$$H = \ln(S) - 12 \cdot \sigma^2$$

where:

- H is the Shannon diversity index (Shannon 1948)
- S is the number of species in the population universe
- σ is the logarithmic standard deviation.

For simplicity, we use ‘sequences’ instead of ‘species’ to refer to the different populations that are members of the community, including species and their microdiversity.

For a different number of sequences (S) and for each σ (from 0.1 to 2.5), we compared the value of H calculated with the formula approach versus the average of H calculated from 1000 random microbial communities designed following a lognormal distribution with standard deviation σ and μ = 0. To do this, we subset S values from a lognormal distribution with standard deviation σ and μ = 0 and normalize these values by the total number of counts. Then, we did 1,000,000 weighted random samplings of these sequences to determine the abundance of each sequence in a sample with 1,000,000 reads. Thus, we generate a community of S sequences and 1,000,000 of reads (total counts). We did 1000 replicas for each pair of values and calculate the average Shannon index of these communities. Finally, we fit a degree-2 polynomial regression by the ordinary least square (OLS) (functions *‘ols’ and ‘fit’* from the ‘statsmodels’ python package) to validate the adjust.

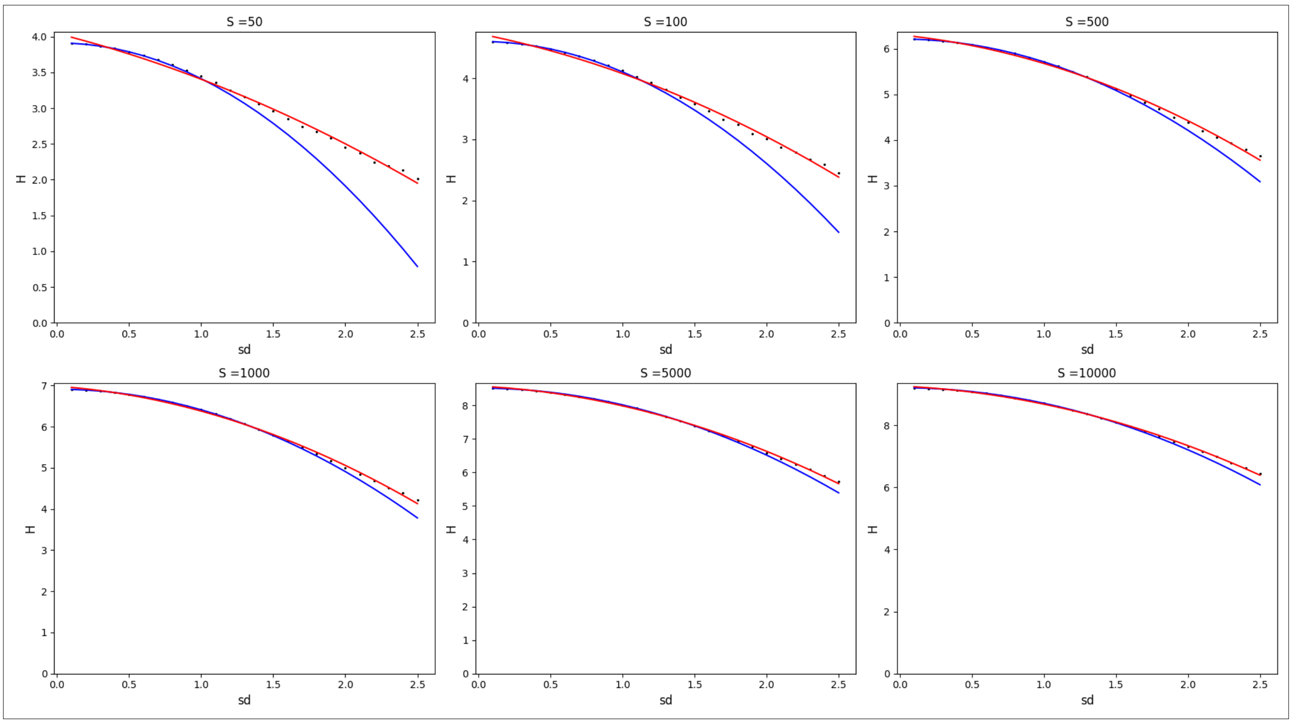


**Fig. S3**: **H depends on σ.** Each graph corresponds to a different number of sequences. X-axis shows the different values of σ, y-axis shows the H calculated for each σ. The blue line represents H calculated with the formula approximation. The average H of the 1000 replicates are indicated with black points, the adjustment to a degree-2 polynomial regression is depicted with a red line. All the regressions were significant $p < 0.05$ (Test F-Fischer).

We observe that for complex communities with a higher number of sequences, H correlates with the σ of the lognormal distribution (**Fig S3**). Therefore, we approximate σ using a given user H clearing in the Longuet and Higgins equation:

$$\sigma = \sqrt[2]{-2 \cdot \left(H - \ln\left(S\right)\right)}$$

Note that in our case, as it has been previously mentioned, S refers to the number of unique sequences that are in the sample: each SILVA sequence which is considered to be a species is divided into unique sequences that are treated as the microdiversity within that species. Consequently, H is referred to the Shannon index calculated at the microdiversity level, instead of a species level. We have also added a script to calculate H at different rank levels so that the user knows which is the resulting community's H.

We allow the user to select S and H as input parameters, but both parameters are connected so note that not all the combinations are possible: if there is only one species in the sample, H must be 0. Additionally, the formula cannot be applied if $\ln(S) < H$ given that $H-\ln(S) <0$ and it is not possible to calculate its square root in the set of all real numbers.

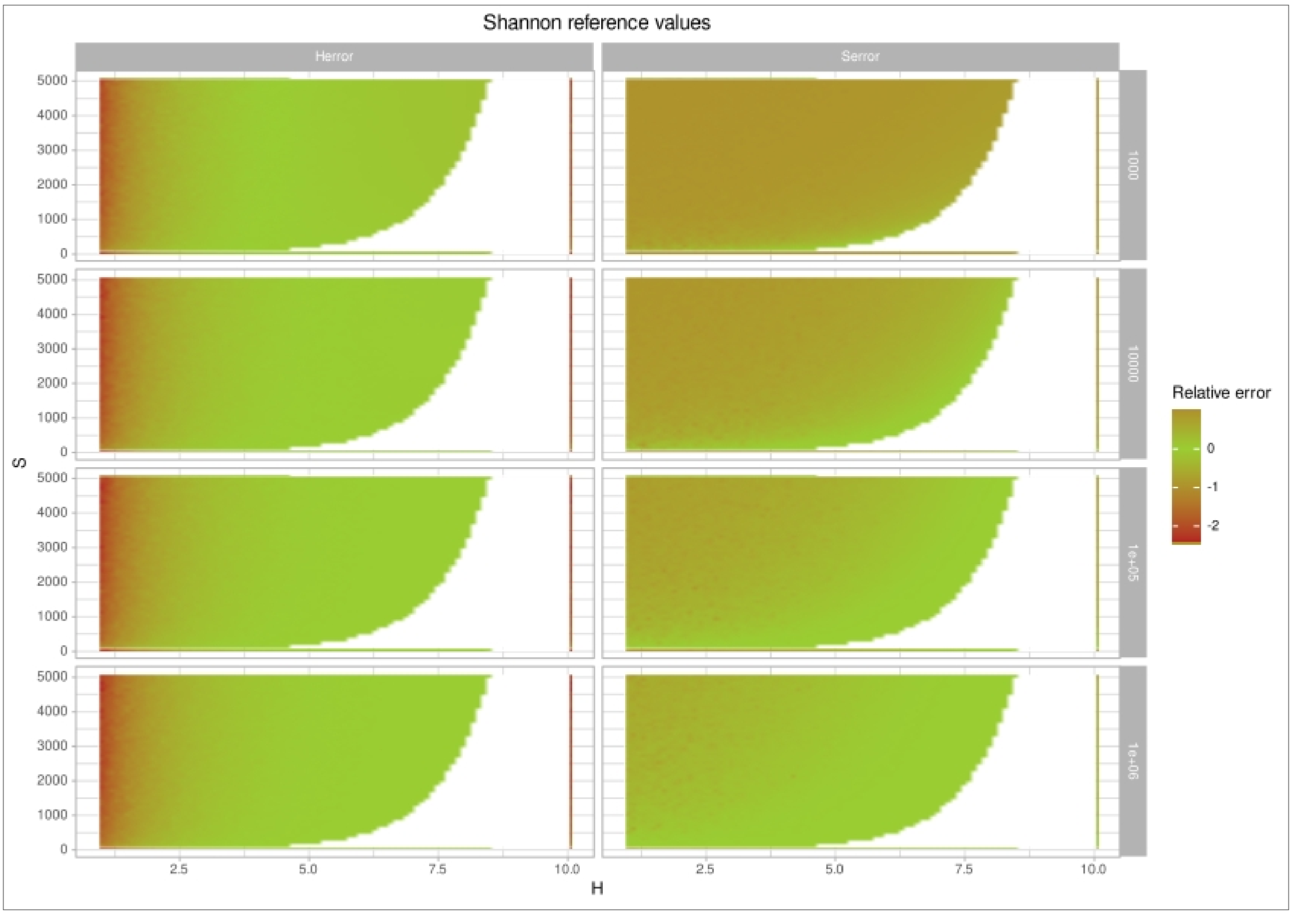


**Fig. S4. Relative error of M&Ms according to the number of reads (rows) when generating the mock with a requested values of S and H.** Green areas are those with a relative error close to 0. Red areas are those with higher relative errors. White area are those values which are not possible according to the equation.

In order to guide the user to select the combinations of values of S, H and number of reads, we have provided **Fig. S5** as a reference. For different values of S and H, we have calculated the minimum number of reads that is required to obtain a mock community with an H and S values with a relative error lower than 5%. Gray area shows those combinations of values that are not possible according to the equation.

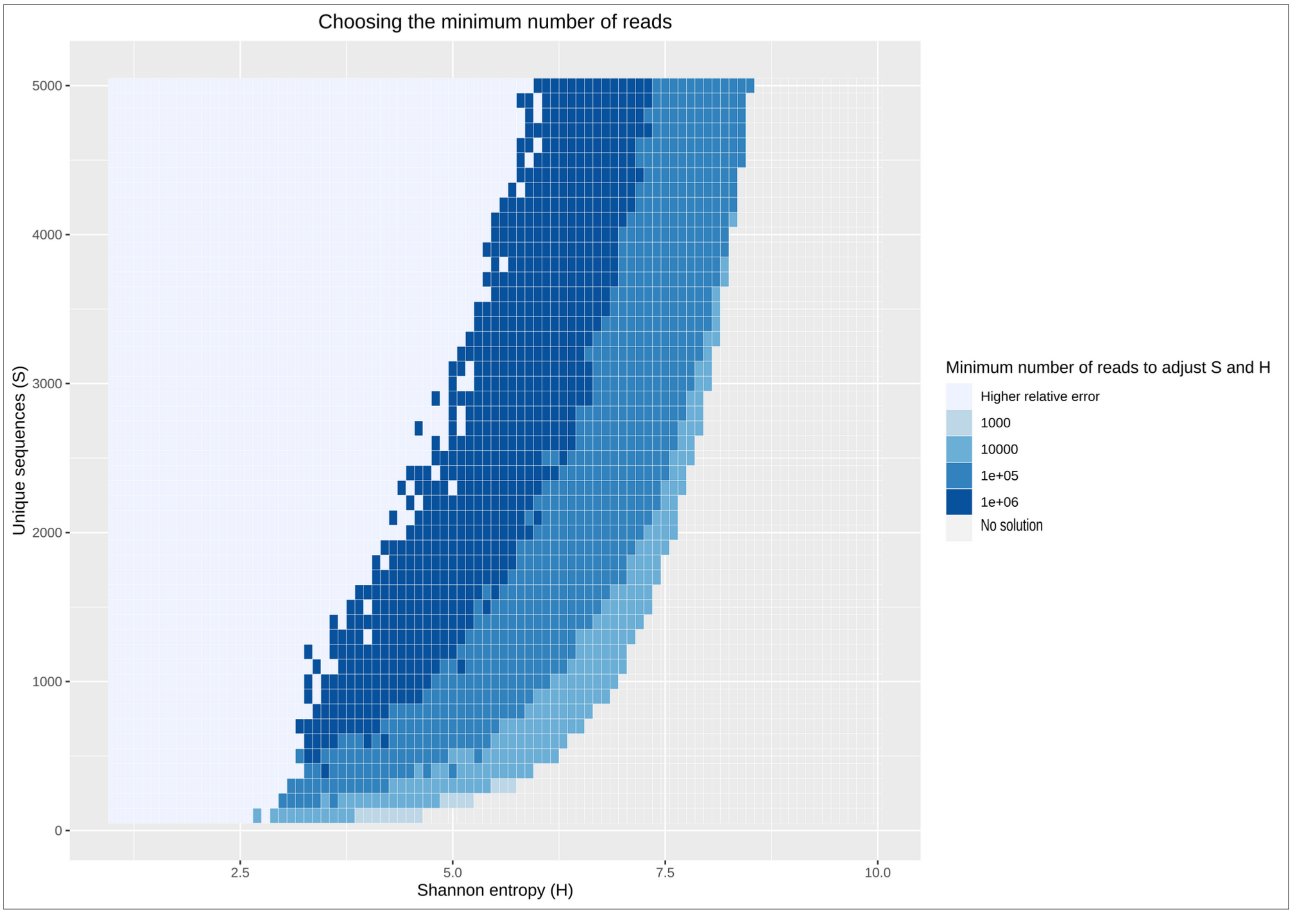


**Fig. S5. Guide to select values of Shannon entropy (H), unique sequences (S) and reads.** Minimum number of reads required to obtain a mock community with a final value of S and H with a relative error less than 5% of the requested values.

### Note 3: Comparison with other simulation tools

We regard that there are two different types of tools. On the one hand, there are tools focused on generating simulated DNA data, e.g. ART (Huang *et al.*, 2012), Grinder (Angly *et al.*, 2012), InSilicoSeqs (Gourlé *et al.*, 2019), MetaSim (Richter *et al.*, 2008) or NanoSim (Yang *et al.*, 2017); some of these tools includes the option to choose different abundance distributions such as the lognormal, exponential or Zero Inflated lognormal distributions. However, these tools have not been designed to simulate realistic microbial communities, but to mimic read-level artifacts in the different sequencing platforms. In such cases, the user has to select the sequences and an abundance distribution model. There is a vast amount of these tools, and, as we have indicated in the main text, the review of Alosaimi *et al.*, 2020, gathers some of them. We compare M&Ms with few examples in **Table S2**. On the other hand, other tools are addressed to simulate realistic species abundance tables but without emulating sequencing data, such as metaSPARSIM (Patuzzi *et al.*, 2019), Community Simulator (Marsland *et al.*, 2020) or SPIEC-EASI (Kurtz *et al.*, 2015). We have included some of these tools in **Table S2**.

M&Ms combines both approaches: it generates artificial microbial community profiles and integrates the InSilicoSeqs software to simulate sequencing. Additionally, M&M's produces a FASTA file and an abundance file, which can be used as inputs for sequencing read simulators other than InSilicoSeqs. To our knowledge, the first tool that also joins both aspects is CAMISIM (Fritz *et al.*, 2019) which automatically produces metagenomic samples emulating different microbial abundance profiles, multi-sample time series, differential abundance studies and strain-level diversity. However although CAMISIM is a very complete tool, M&Ms incorporates most of these functions, but also allows users to design realistic multi-sample mock communities of known inter-taxa correlations and based on ecologically meaningful parameters (source environment, number of species, Shannon diversity).

| | | | Reads simulators | | Microbial Community Simulators | | |
|---|---|---|---|---|---|---|---|
| | M&Ms | CAMISIM | ART | Grinder | metaSPARSIM | Community Simulator | SPIEC-EASI |
| Year | 2020 | 2019 | 2012 | 2012 | 2019 | 2020 | 2015 |
| Last updated | 2020 | 2020 | 2016 | 2016 | 2020 | 2020 | 2020 |
| Reads generation | Yes | Yes | Yes | Yes | No | No | No |
| Microdiversity simulation | Yes | Yes | No | No | No | No | No |
| Requires a real community | No | No | No | No | No | Yes | Yes |
| Abundance profile | Yes: LN | Yes: LN | No | Yes: U, L, PL, LN, EX | Yes: GM | Yes (resource concentrations) | Yes: previous community |
| Abundance table | Yes | Yes | No | No | Yes | Yes | Yes |
| Sampling replicates | No | Yes | No | No | Yes | No | No |
| Species correlation | Yes: ZINBD | Yes: Time series | No | No | Yes: MHG | Yes (resource concentrations) | Yes: ZINBD, ZIP |
| Environment | Yes | No | No | No | No | No | Yes |
| Shannon Index (H) | Yes | No | No | No | No | No | No |
| Richness (α- diversity) | Yes | Yes | Yes | Yes | Yes | Yes | Yes |

**Table S2. Comparison of M&Ms with similar tools.** LN: lognormal distribution, ZINBD: Zero-Inflated Negative Binomial Distribution, U: Uniform distribution, L: Linear distribution, PL: powerlaw distribution, LN: logarithmic distribution, EX: Exponential distribution, GM: Gamma distribution, MHG: Multivariate Hypergeometric distribution, ZIP: Zero Inflated Poisson Distribution.
